# Dysfunctional TRPM8 signalling in the vascular response to environmental cold in ageing

**DOI:** 10.1101/2021.05.10.443379

**Authors:** Dibesh Thapa, João de Sousa Valente, Brentton Barrett, Fulye Argunhan, Sheng Y. Lee, Sofya Nikitochkina, Xenia Kodji, Susan D. Brain

**Affiliations:** Section of Vascular Biology and Inflammation, School of Cardiovascular Medicine and Sciences, BHF Centre of Research Excellence, King’s College London, Franklin-Wilkins Building, Waterloo Campus, King’s College London, London SE1 9NH, UK; Cancer Research UK, Cambridge Institute, University of Cambridge, Cambridge, CB2 0RE, UK; Skin Research Institute, Agency of Science, Technology, and Research (A*STAR), 8A Biomedical Grove, #06-21 Immunos, Singapore 138648

## Abstract

Ageing is associated with increased vulnerability to environmental cold exposure. Previously, we identified the role of the cold-sensitive transient receptor potential (TRP) A1, M8 receptors as vascular cold sensors in mouse skin. We hypothesised that this dynamic cold-sensor system may become dysfunctional in ageing. We show that behavioural and vascular responses to skin local environmental cooling are impaired with even moderate ageing, with reduced TRPM8 gene/protein expression especially. Pharmacological blockade of the residual TRPA1/TRPM8 component substantially diminished the response in aged, compared with young mice. This implies the reliance of the already reduced cold-induced vascular response in ageing mice on remaining TRP receptor activity. Moreover, sympathetic-induced vasoconstriction was reduced with downregulation of the α_2c_ adrenoceptor receptor in ageing. The cold-induced vascular response is important for sensing cold and retaining body heat and health. These findings reveal that cold sensors, essential for this neurovascular pathway, decline as ageing onsets.

## Introduction

Upon exposure to cold, depending on the type and intensity, several counterbalancing responses are produced, such as the behavioural response of shivering thermogenesis involving skeletal muscle, or biochemical responses such as non-shivering thermogenesis in brown adipose tissue (BAT) and peripheral vasoconstriction in skin (Señarís et al., 2018, Morrison, Shaun F., Nakamura, 2011, Morrison, S. F., Nakamura, 2019). To produce such responses, thermo-sensors in the form of temperature sensitive sensory receptors are distributed throughout the skin and are considered to work as a first line of defence against cold, which makes peripheral cutaneous responses a fundamental event in the defence against environmental thermal challenge. The sensory receptors in the skin initiate the vascular cold constrictor response which acts to protect against body heat loss and prevent hypothermia. This response is followed by the subsequent vasodilation, a restorative response that is essential to protect the affected skin against cold-induced conditions, such as chilblains, trench foot, frostbite, and Raynaud’s condition (Daanen, van der Struijs, Norbert R., 2005, Keatinge, 1957, Lewis, 1930). It is a finely tuned well balanced response that maintains cellular function and physiological homeostasis during cold exposure. Whilst this response is relevant to all ages, physiological changes in ageing leads to dysfunctional signalling which causes a reduced adaptation to cold exposure (Guergova, Dufour, 2011). With the lack of physical activity in the elderly population, it exacerbates the fall in core body temperature which can cause fatal cardiovascular and respiratory problems (Billeter et al., 2014, Stares, Kosatsky, 2015). This is normally the biggest cause behind the NHS excess winter deaths that we witness every year, where in 2018 it caused approximately 11,000 deaths linked to cold exposure in England (Office for National Statistics, 2019).

We have previously delineated the primary roles of transient receptor potential (TRP) channels in producing a distinctive biphasic vascular response to cold in the mouse paw consisting of a TRP ankyrin 1 (TRPA1)/ melastatin 8 (TRPM8)-initiated sympathetic α_2c_ adrenoceptor mediated neuronal vasoconstriction and a distinct TRPA1-CGRP mediated sensory-vasodilator component (Aubdool et al., 2016). TRPA1 is a biomolecular sensor for noxious cold (<18°C), mediating aversive behaviour such as avoiding cold-induced pain, whilst also being involved in mediating inflammatory pain (Kwan et al., 2006, Nassini et al., 2014, Jain et al., 2011, Gouin et al., 2017). Additionally, it activates C and Aδ sensory nerves to release neuropeptides such as CGRP to mediate neurogenic vasodilation (Aubdool et al., 2016, Story et al., 2003, Gentry et al., 2010). TRPM8 is sensitive to cool temperatures (<28°C) (McKemy, Neuhausser & Julius, 2002, Peier et al., 2002). It is involved in deep body cooling and suggested to supersede the role of TRPA1 (Gavva et al., 2012). TRPM8 is also suggested to be a vasoactive stimulus (Bautista et al., 2007, Johnson et al., 2005, Silva et al., 2019). The other established receptor that plays a pivotal role in cold signalling is the sympathetic α_2c_ adrenoceptor, which mediates the vasoconstriction of the blood vessels (Bailey et al., 2004). Whilst the sympathetic branch that is involved in the vasoconstrictor component of the cold response has been shown to have reduced activity in ageing humans (Holowatz, Thompson-Torgerson & Kenney, 2010, Degroot, Kenney, 2007), little is known about the functionality of the cold receptors TRPA1 and TRPM8 in ageing. In the current study we hypothesize that signalling via the cold receptors TRPA1 and TRPM8 deteriorates with ageing which causes an impaired vascular response to the cold.

The primary objective of this study is to investigate the cutaneous vascular response to cold in ageing, focusing on the activity of cold TRP receptors; TRPA1 and TRPM8. As sympathetic-sensory neuronal signalling is key for the cutaneous vascular cold response in ageing, we also searched for evidence of dysfunction within these systems. Here using *in vivo*, *ex vivo*, genetic, and pharmacological approaches we show that TRPA1 and TRPM8 signalling declines with ageing which affects the sensing as well as functional pathways involved in cold signalling; all of which contribute to the impaired cold vascular response. Additionally, we provide evidence that the α_2c_ adrenoceptor as well as the TRPM8 receptor both play critical roles to influence this outcome, as the expression of both diminishes significantly in ageing which impacts the vascular response to cold. These important findings establish the dynamic role of cold sensitive TRP receptors and sympathetic receptors in the cutaneous vascular response to the cold as ageing occurs.

## Results

### Cold-induced vascular response is impaired in ageing

We analysed the cold induced vascular response in WT CD1 females (Young: 2-3 months, Aged: 13-15 months) with full-field laser speckle imager (FLPI) using the cold water immersion model (Fig 1a) developed in our laboratory (Aubdool et al., 2014, Pan et al., 2018). After the baseline blood flow was measured for 5 min, the ipsilateral hindpaw was immersed in cold water at 4°C, a temperature that produces a robust vascular response, for 5 min and blood flow was then recorded for another 30 min. The cold treatment produced a typical vascular response of rapid vasoconstriction followed by a prolonged recovery vasodilator response in both young and aged mice (Fig 1b-c, Supplementary Fig 1a). In young mice, the cold treatment produced a maximum vasoconstriction of 51.1 ± 1.176%, however, in aged mice this was significantly blunted with maximum vasoconstriction of 27.7 ± 2.976% (Fig 1d). These changes were reflected in the area under the response curve (AUC) analysis with a significantly greater response in young than aged mice (Fig 1e). The result was extended by measurement of the blood flow recovery after the cold treatment. Although blood flow did not fully recover back to the baseline, the initial rate of recovery immediately after maximum vasoconstriction before it slowly plateaued off was significantly faster in the young mice compared to the aged mice (Fig 1f). These results suggest that with ageing the cold induced vascular response starts to diminish, which affects both parts of the vascular response. We were surprized that these changes were observed with moderately aged mice, equivalent to middle aged in human terms (Dutta, Sengupta, 2016). However, at this age there is a clear evidence of elevated gene expression in DRG and skin of senescence markers associated with ageing, p16 and p21, (Fig 1g-h) also confirmed by western blotting (Fig 1i).

**Figure 1:**
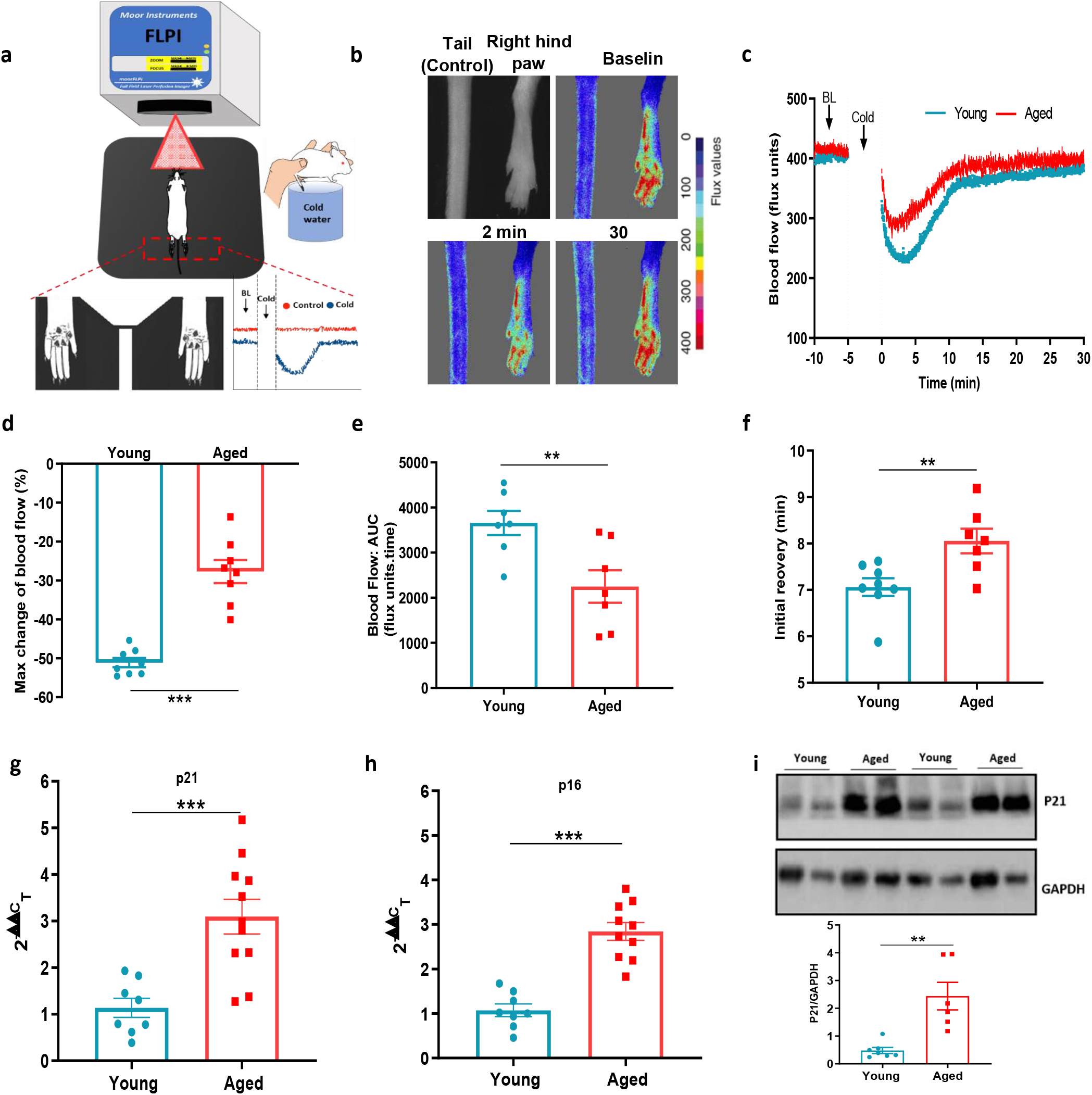
Cold-induced vascular response is impaired with ageing. **(a)** Diagram illustrates the experimental setup of cold-induced vascular response protocol; FLPI from top measures the blood flow in the hindpaw of the anaesthetized mouse when on a heating mat in response to cold water immersion. The expanded component (highlighted by dotted red lines) shows the hindpaw region in which the blood flow is recorded, and a graph of typical blood flow response is shown. Recording is paused for cold treatment where one of the hindpaw is immersed in cold water for 5 min. **(b)** Representative FLPI image shows the blood flow in cold-treated hind paw at baseline, 2 min and 30 min after the cold water treatment. **(c)** Graph shows the raw blood flow trace (mean) of vascular response with cold (4°C) water treatment (n=8). **(d)** % change in hindpaw blood flow from baseline to 0-2min following cold water treatment (maximum vasoconstriction). **(e)** The AUC to maximum vasoconstriction point assessed by area under the curve (AUC). **(f)** Time of blood flow recovery immediately after maximum vasoconstriction until the start of the plateau period. **(g-h)** RT-PCR CT analysis shows fold change of p21 and p16 gene expression normalized to three housekeeping genes in dorsal root ganglia (DRG) of young and aged mice. **(i)** Representative western blot of p21 in hindpaw skin of young and aged mice and densitometric analysis normalized to GAPDH. (BL = baseline). Data is presented as mean and all error bars indicate s.e.m. **p<0.01, ***p<0.001. (Two-tailed Student’s t-test).

To extend our mechanistic understanding, we also used a laser Doppler imager (VMS-LDF), in addition to FLPI, which simultaneously measures the blood flow, skin temperature and tissue oxygen saturation level at a single point, to investigate the vascular response to cold. Similar to the results obtained using the FLPI, the environmental cold water treatment produced an impaired vascular response in the paws of aged mice compared to the young mice (Fig 2a). In young mice, the cold treatment produced a maximum vasoconstriction of 45.5 ± 2.952%, however in aged mice this was significantly lower with a maximum vasoconstriction of 23.4 ± 4.678%, a result which was reflected in AUC analysis (Fig 2b-c). There was a trend of greater reduction in skin temperature of aged mice after the cold water treatment; however, the aged mice had a significantly higher skin temperature at baseline, suggesting they were losing more body heat and consistent with the fact that the ability to maintain core body temperature declines with ageing (Fig 2d-f). The tissue oxygen saturation level underwent a similar reduction in both young and aged mice after the cold exposure (Fig 2g-h) but recovered more robustly in the young mice compared to the aged mice as shown by AUC analysis (Fig 2i). We also found evidence of increased cellular stress as protein expression of 3-nitrotyrosine, a biomarker of oxidative stress produced via reactive nitrogen species was elevated in aged hindpaw skin (Supplementary Fig 2), in keeping with physiological ageing. These results from two distinct techniques confirm our finding that the cold induced vascular response starts to diminish with ageing.

**Figure 2:**
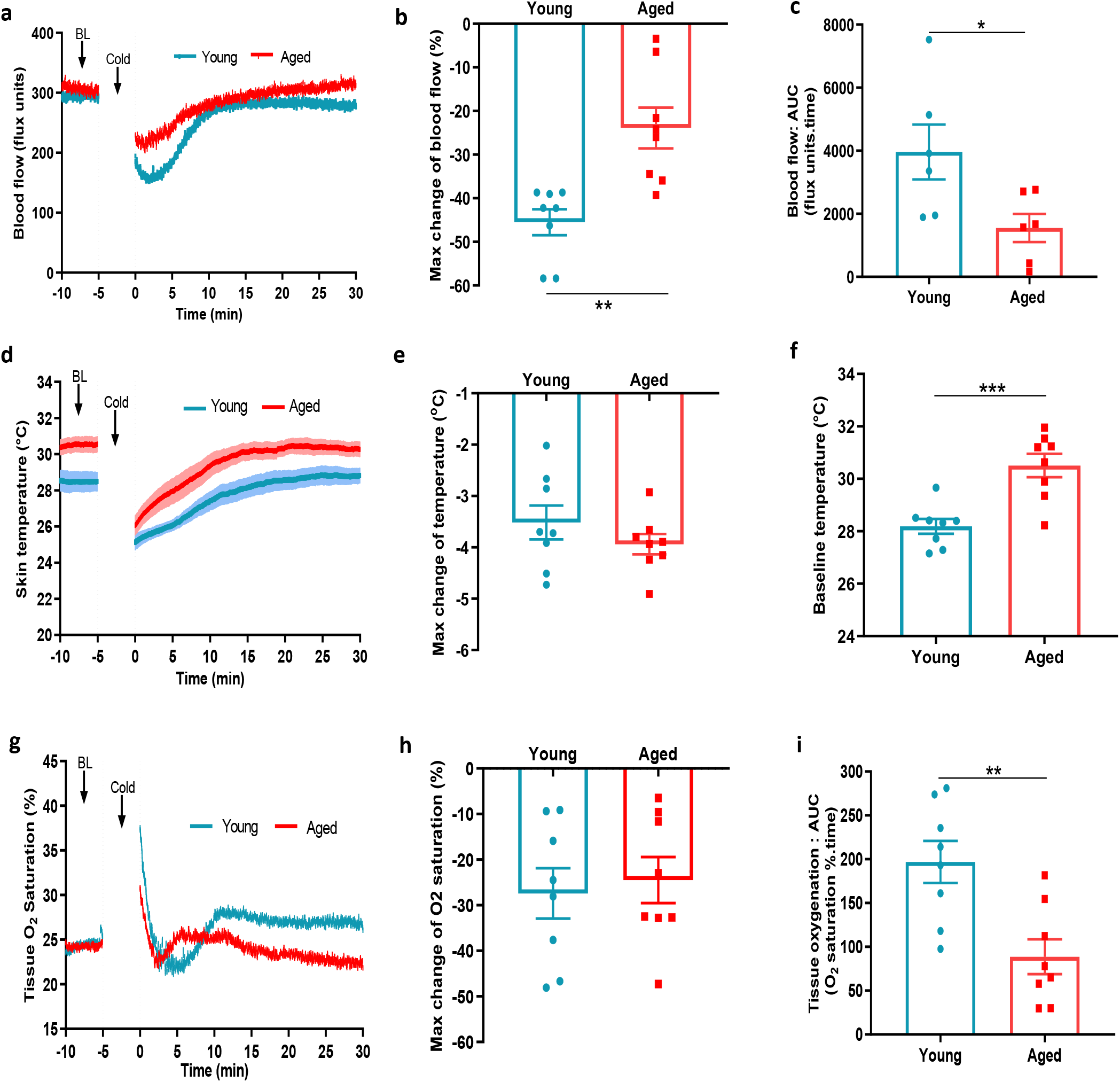
Blood flow, skin temperature and tissue oxygen saturation with cold treatment in ageing. **(a)** Mean blood flow trace of the vascular response with cold (4°C) water treatment (n=8). **(b)** % change in hindpaw blood flow from baseline to 0-2min following cold treatment (maximum vasoconstriction). **(c)** The vasoconstriction response caused by cold water treatment represented by area under curve (AUC). **(d)** The mean blood flow (± s.e.m.) recordings of hindpaw skin temperature with cold water treatment. **(e)** Maximum reduction in skin temperature following 5 min cold treatment. **(f)** The baseline skin temperature. **(g) %** mean tissue oxygen saturation during cold water treatment. **(h)** % maximum change in tissue oxygen saturation from baseline following cold water treatment **(i)** % tissue oxygen saturation recovery after cold water treatment assessed by area under the curve. (BL = baseline). Data is presented as mean and all error bars indicate s.e.m. *p<0.05, **p<0.01, ***p<0.001. (Two-tailed Student’s t-test).

### Cold sensitivity is impaired in ageing

To learn if cold sensitivity had altered with ageing, we examined the functionality of TRPA1 and TRPM8 channels in behavioural studies using a cold plate set at 4°C, 10°C, and 20°C, within the activation range of TRPA1 and TRPM8 receptors (Dhaka et al., 2007, Kwan et al., 2006). At all three cool/cold temperatures, the aged mice showed a significant delayed latency for paw licking/paw withdrawal/jumping compared to the young mice, suggesting impaired cold sensing in aged mice (Fig 3a-c), but with little difference in the total number of responses observed among groups (Fig d-f). When the test was performed at 30°C, a temperature outside the activation range of TRPA1 and TRPM8, we observed no delayed latency in response time, although the total number of responses was significantly lower in the aged mice (Fig 3g-h). These results indicate that there is a reduction in sensitivity to cold with ageing at temperatures at which the cold sensors TRPA1 and TRPM8 are active; thus leading us to hypothesise that at least one cold-sensitive TRP pathway deteriorates with ageing. Of note, the largest difference in response time between young and aged mice was observed at 20°C, in keeping with the TRPM8 activation range (Fig 3i).

**Figure 3:**
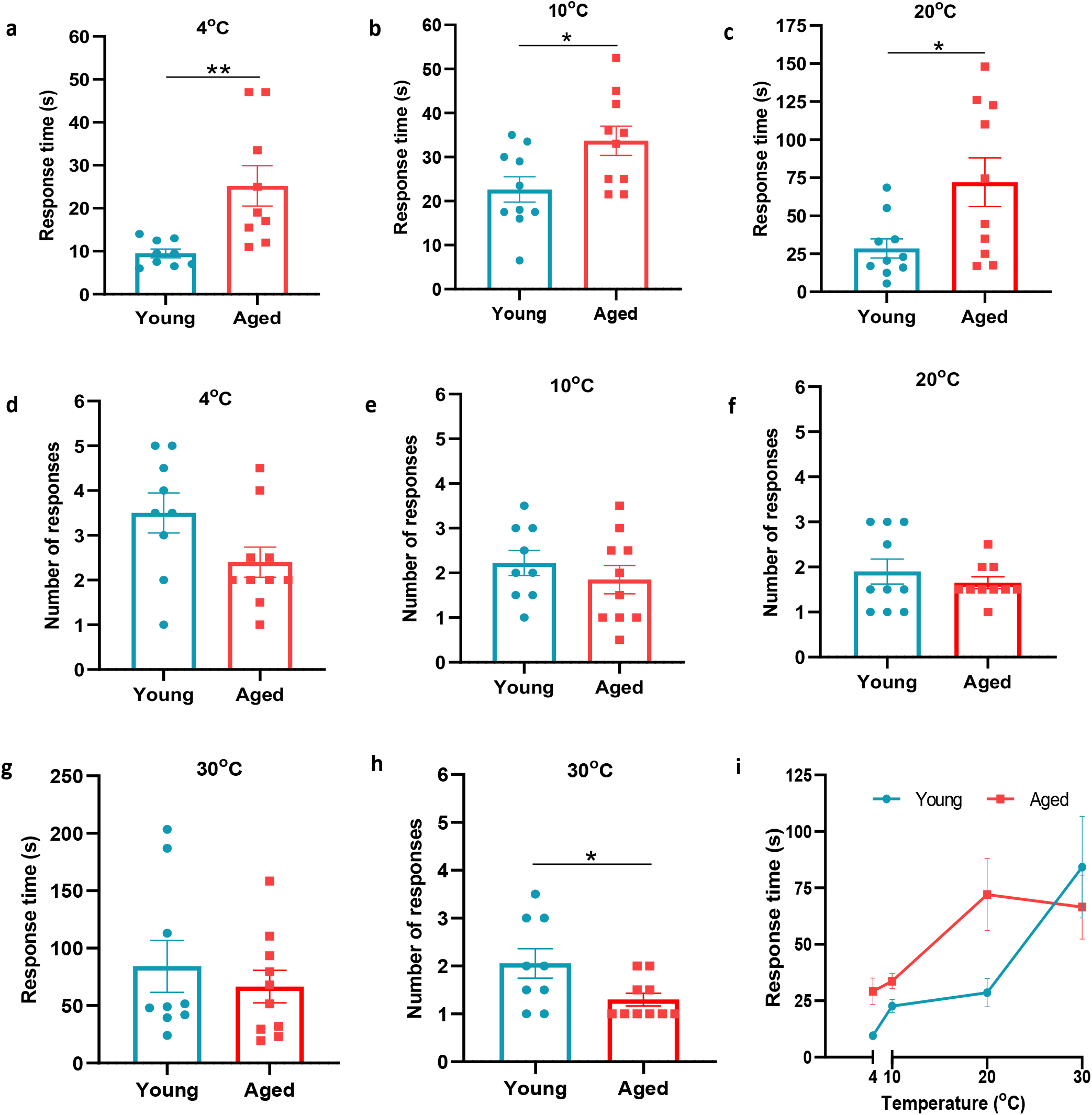
Behavioural analysis with cold plate in young and aged mice. **(a-c)** Time of first response of mice to cold plate set at 4°C, 10°C, and 20°C. **(d-f)** The total number of responses from mice during the cold plate experiment at 4°C, 10°C, and 20°C. **(g-h)** Time of first response of mice to cold plate set at 30°C and total number of responses. **(i)** Line graph illustrates the difference in mean response time at the four different temperatures the cold plate assay was performed, between young and aged mice. All results are shown as mean ± s.e.m. *p<0.05, **p<0.01. (Two-tailed Student’s t-test).

### The cold-induced vascular response remains dependent on TRPA1 but not TRPM8 in ageing

To investigate the role of TRPA1 and TRPM8 in the local cold water immersion test; we measured the cold-induced vascular response in the presence of the TRPA1 antagonist A967079 (100 mg kg^−1^ i.p.) and TRPM8 antagonist AMTB (10 mg kg^−1^ i.p.), a combination previously shown by us to inhibit the cold induced vascular response (Pan et al., 2018). The combined pre-treatment of A967079 and AMTB partially but significantly inhibited the vasoconstriction in young mice. By comparison, this treatment regime produced a more substantial inhibition of vascular responses induced by cold in the aged mice (Fig 4a, 4d). This result reveals that the role of TRP receptors in the cold-induced vascular response remains and suggests as ageing occurs the TRP-mediated signalling may become more important. Next, we performed the cold water immersion test in the presence of either A967079 or AMTB. The A967079 treatment produced a similar effect to the combined antagonist treatment of A967079+AMTB, where the antagonist was more effective in aged than in the young mice (Fig 4b, Fig 4e). By comparison, the AMTB treatment inhibited the response in young mice, but had no significant effect in the vascular response to cold (Fig 4c, 4f) in aged mice. This provides further evidence that as ageing occurs, TRPM8 loses its ability to respond to local cold treatment. Next, we examined the expression of TRPA1 and TRPM8 in DRGs of young and aged mice. RT-PCR analysis of DRG showed similar level of TRPA1 mRNA in both young and aged mice (Fig 4g). However, the level of TRPM8 mRNA was significantly reduced in the aged compared to the young mice (Fig 4h), as was its protein expression when analysed by western blot (Fig 4i).

**Figure 4:**
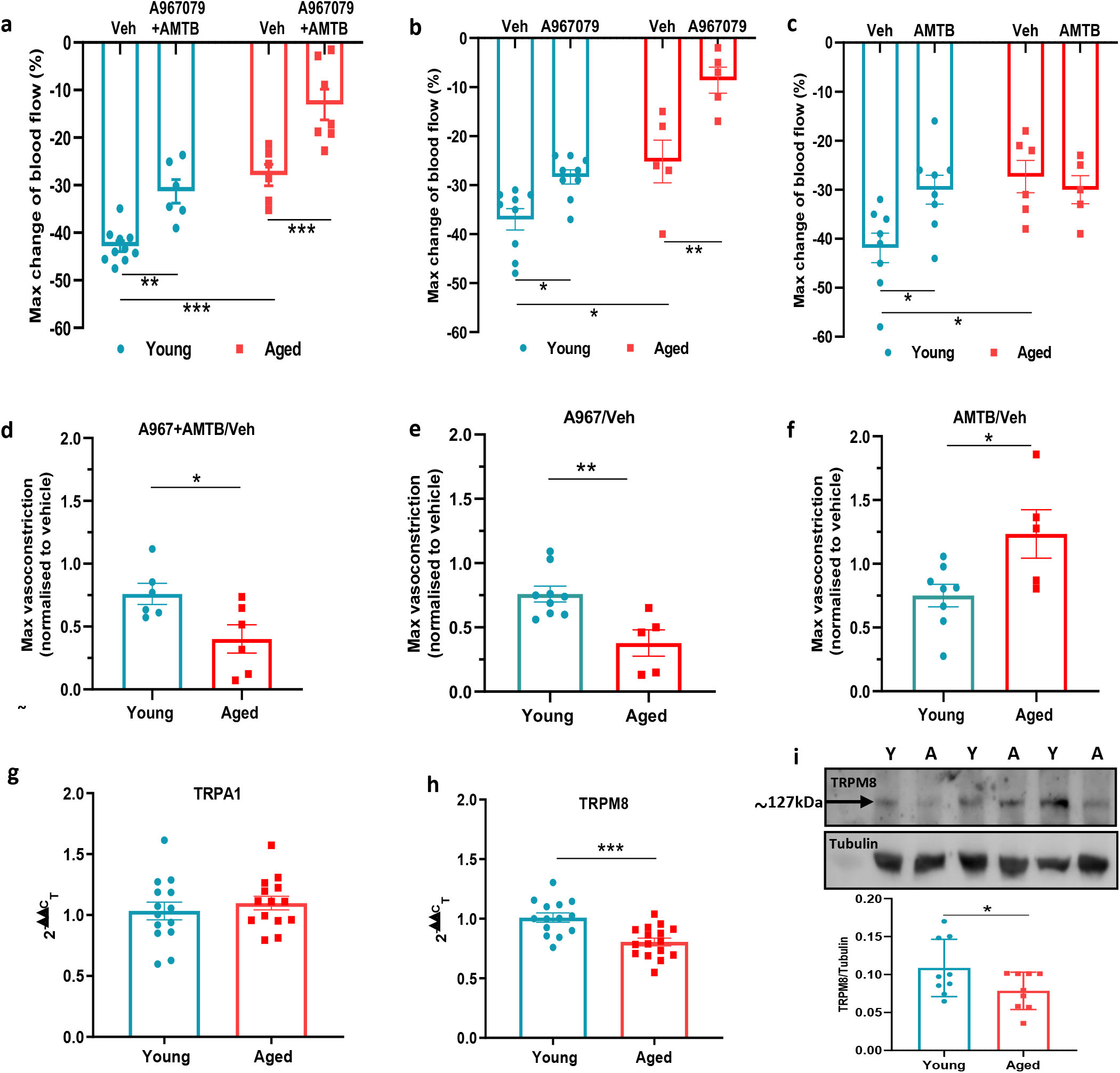
TRPA1 and TRPM8 are involved in cold-induced vascular response. Vascular responses with cold (4°C) water treatment in mice pre-treated with combined TRPA1 antagonist A967079 (100 mg kg^−1^) and TRPM8 antagonist AMTB (10 mg kg^−1^), or vehicle control (Veh - 10% DMSO, 10% Tween in saline) i.p. 30 min before cold treatment. **(a-c)** % change in hindpaw blood flow from baseline to 0-2min following cold treatment (maximum vasoconstriction) in mice treated with combined antagonist **(a)** A967079+AMTB, **(b)** A967079, and **(c)** AMTB. **(d-f)** Maximum vasoconstriction caused by cold water treatment in mice treated with combined antagonist **(d)** A967079+AMTB, **(e)** A967079, and **(f)** AMTB normalized against vehicle treated mice. **(g-h)** RT-PCR CT analysis shows fold change of **(g)** TRPA1 and **(h)** TRPM8 normalized to three housekeeping genes in dorsal root ganglia (DRG). **(i)** Representative western blot of TRPM8 in DRG of young and aged mice and densitometric analysis normalized to Tubulin (Y=young, A=aged). All results are shown as mean ± s.e.m. *p<0.05, **p<0.01, ***p<0.001. (Two-way ANOVA with Tukey’s *post hoc* test or Student’s t-test).

### TRPA1 and TRPM8 vasodilator signalling is impaired in ageing

Thus far we had gained multiple evidence that TRPM8 activity is impaired in vascular signalling in ageing, with some evidence for a reduction in TRPA1 activity. To build on these findings, we examined the vasoactive effect of TRPA1 and TRPM8 agonists that are commonly associated with sensory nerves. The topical application of cinnamaldehyde (CA) and menthol on mouse skin have previously been shown to mediate vasodilation via TRPA1 and TRPM8 channels respectively (Craighead et al., 2017, Aubdool et al., 2016). The topical application of menthol (10%) to the ear caused increased blood flow in young mice, which was significantly lower in the aged mice (Fig 5a), as shown by the maximum increase in blood flow (Fig 5b). The AUC analysis of blood flow showed significant increase with menthol treatment compared to vehicle in young mice but not in aged mice (Fig 5c). Similarly, cinnamaldehyde (CA, 10%) application also increased blood flow in young mice, however, this increase was significantly lower in aged mice (Fig 5d-e, Supplementary Fig 3a-b). The AUC analysis showed a significant increase in blood flow with CA treatment compared to vehicle in young mice but not in aged mice (Fig 5f). These findings suggest that the TRPA1 and TRPM8-mediated vasoactive activity starts to deteriorate in moderate ageing, and it is not exclusive to cold signalling. To build on this concept, we examined whether the activity of another prominent TRP receptor, TRPV1, is also impaired with ageing. To probe this, we studied capsaicin-induced increase in ear blood flow (Grant et al., 2005). The topical application of 10% capsaicin produced a similar increase in ear blood flow in both young and aged mice (Fig 5g-i) indicating, unlike the TRPA1 and TRPM8 signalling, the TRPV1 signalling does not deteriorate with ageing.

**Figure 5:**
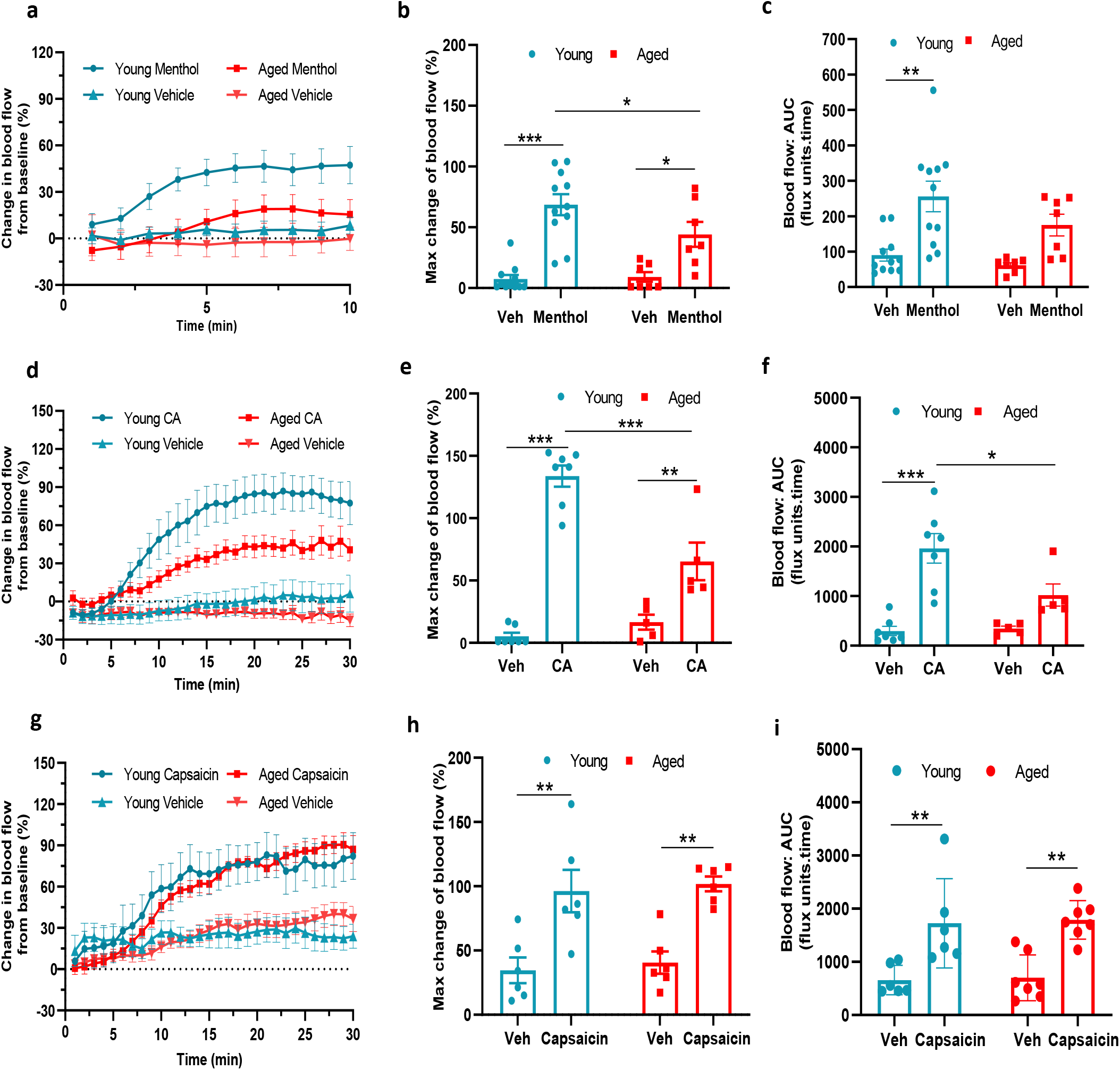
TRPA1 and TRPM8 activity deteriorates with ageing. **(a)** Graph shows the % mean ± s.e.m. of blood flow change from baseline in response to topical application of menthol (10%) and vehicle (Veh - 10% DMSO in ethanol) in ear of young and aged mice. **(b)** % maximum change in ear blood flow induced by menthol application in young and aged mice. **(c)** AUC analysis of % blood flow increase from baseline after menthol application compared to vehicle. **(d)** Graph shows the % mean ± s.e.m. of blood flow change from baseline in response to topical application of cinnamaldehyde (10% CA) and vehicle (10% DMSO in ethanol) in ear of young and aged mice. **(e)** % maximum change in ear blood flow induced by CA application in young and aged mice. **(f)** AUC analysis of % blood flow increase from baseline after CA application compared to vehicle. **(g)** Graph shows the % mean ± s.e.m. of blood flow change from baseline in response to topical application of capsaicin (10%) and vehicle (10% DMSO in ethanol) in ear of young and aged mice. **(h)** % maximum change in ear blood flow induced by capsaicin application in young and aged mice. **(i)** AUC analysis of % blood flow increase from baseline after capsaicin application compared to vehicle. All results are shown as mean ± s.e.m. *p<0.05, **p<0.01, ***p<0.001. (Two-way ANOVA with Tukey’s *post hoc* test).

### Dysfunction in sympathetic signalling contributes to impaired cold response in ageing

In comparison to the sensory system, the importance of sympathetic nerves in mediating the vascular smooth muscle constriction in the cold response is well established (Bailey et al., 2004, Smith et al., 2004). To understand whether there is modulation of this pathway as ageing progresses, we examined the sympathetic-mediated vasoconstriction. The response to the intraplantar injection of the non-selective and endogenous sympathetic neurotransmitter noradrenaline (NA) revealed a significantly greater reduction of blood flow in young mice compared to aged mice (Fig 6a-b). Knowing that the α_2c_ adrenoceptor is essential for cold induced vasoconstriction (Aubdool et al., 2014, Bailey et al., 2004, Honda et al., 2007), we then proceeded to investigate the effect of the selective α_2_ adrenoceptor agonist medetomidine, in hindpaw blood flow. Medetomidine caused immediate vasoconstriction as expected, but the response was blunted in aged mice compared to young mice (Fig 6c-d). These results recapitulate previous findings that suggest a defect also in sympathetic signalling in aged mice involving the α_2c_ adrenoceptor, in addition to the cold TRP receptors. The western blotting analysis of the hind paw skin showed a significant reduction in the expression of α_2c_ adrenoceptor in aged mice (Fig 6e). To elucidate further potential defects in the sympathetic pathway with ageing, we investigated the biosynthesis pathway of NA, the major signalling molecule of sympathetic system. Tyrosine hydroxylase (TH), an enzyme that catalyses the rate limiting stage of noradrenaline synthesis, showed a similar level of expression (Fig 6f) including of its active form, phosphorylated TH in both young and aged mice (Supplementary Fig 4), suggesting the production of noradrenaline remained unaltered with ageing. This indicates that in ageing the expression and function of the α_2c_ adrenoceptor diminishes and that contributes to the impaired constrictor response against cold.

**Figure 6:**
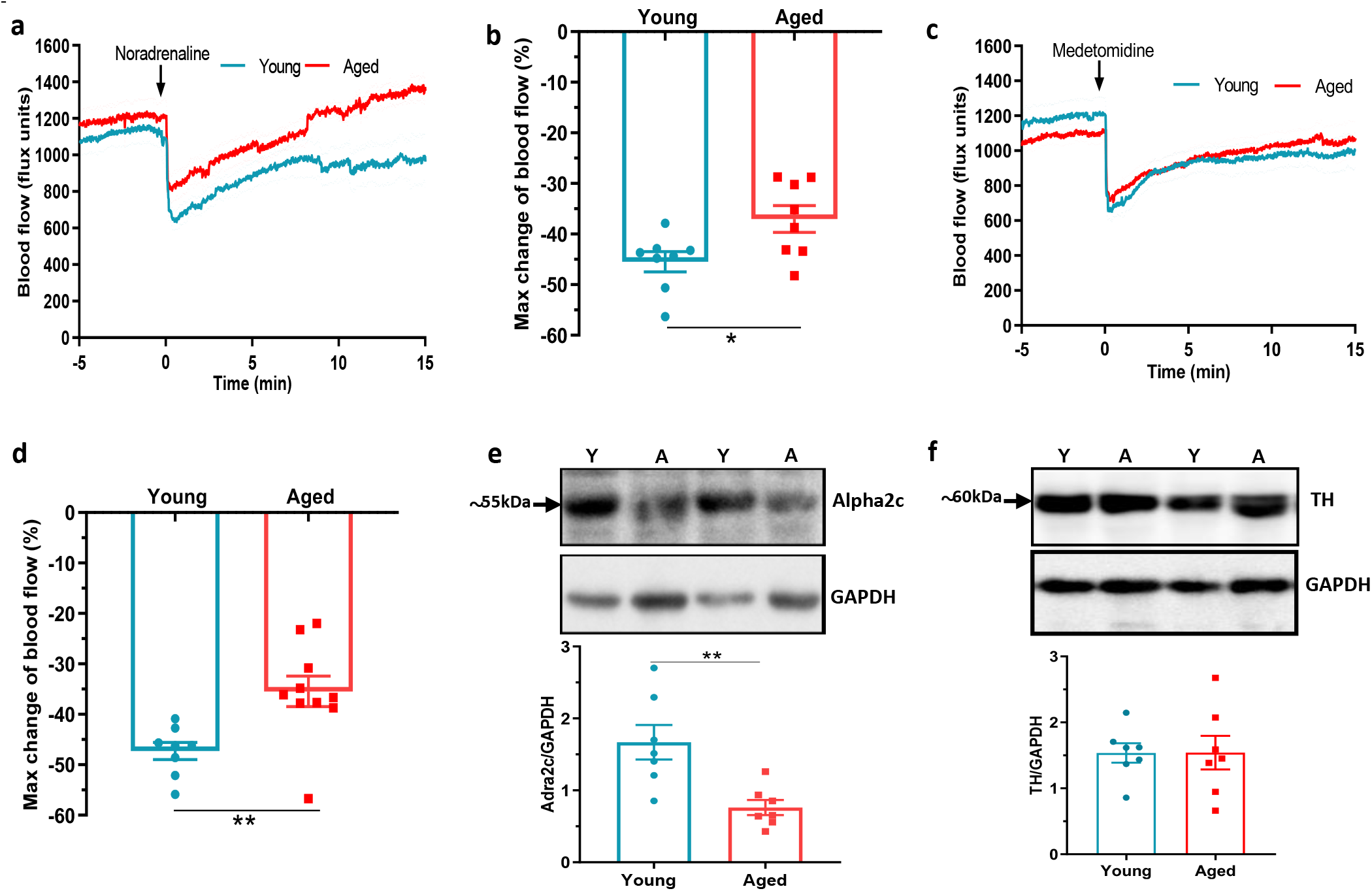
Dysfunctional sympathetic signalling in ageing. **(a)** Graph shows the mean ± s.e.m. blood flow in hindpaw with intraplantar injection of noradrenaline (1.25ng/μl in saline in 20μl) in young and aged mice (n=8). **(b)** % maximum change in blood flow from baseline induced by noradrenaline (maximum vasoconstriction). **(c)** Graph shows the mean ± s.e.m. blood flow in hindpaw with intraplantar injection of medetomidine (1.25ng/μl in saline in 20μl) in young and aged mice (n=8-10) **(d)** % maximum change in blood flow from baseline induced by medetomidine (maximum vasoconstriction). **(e)** Representative western blot of alpha2C (α_2c_) adrenoceptor in mice hindpaw skin with densitometric analysis normalized to GAPDH. **(f)** Representative western blot of tyrosine hydroxylase (TH) in mice hindpaw skin with densitometric analysis normalized to GAPDH (Y=young, A=aged).

### Sympathetic-sensory signalling and influence of ageing

We extended our investigation of sympathetic system in vascular cold response by exploring potential crosstalk between sympathetic and sensory signalling in ageing. To elucidate this, we first investigated DRG and found that α_2a_ and α_2c_ adrenoceptor gene expression was reduced in ageing (Supplementary Fig 5a-b) similar to that shown for the TRPM8 gene, (Fig 4h) whilst no significant difference was found for TRPV1 receptors (Supplementary Fig 5c) in keeping with results for TRPA1 (Fig 4g). By comparison, whilst the TRP receptors are well known to be expressed in sensory neurons there is evidence for a broader localisation (Hirai et al., 2018, Jain et al., 2011, Smith et al., 2004, Yang et al., 2006). We investigated the possible expression of these receptors on sympathetic nerves by collecting the sympathetic ganglia from the cervical and thoracic paravertebral regions where they could directly influence the NA transmission that mediates the vasoconstrictor component of the vascular cold response. To confirm the phenotype of sympathetic neurons, we used positive markers such as tyrosine hydroxylase (TH) and dopamine β-hydroxylase (Supplementary Fig 6) both of which exhibited high expression compared to sensory neuron of DRGs and kidney which were used as negative controls. The RT-PCR data on sympathetic ganglia showed the gene expression of both TRPA1 and TRPM8 in young and aged mice. Interestingly, the expression of both receptors were significantly downregulated in aged mice (Fig 7a-b). Whilst there is no feasible selective TRPA1 antibody available, western blot analysis of TRPM8 on sympathetic ganglia recapitulated the qPCR finding of diminished expression in aged mice compared to young mice (Fig 7c). These findings reveal expression of cold TRP receptors in sympathetic neurons which are diminished in ageing.

**Figure 7:**
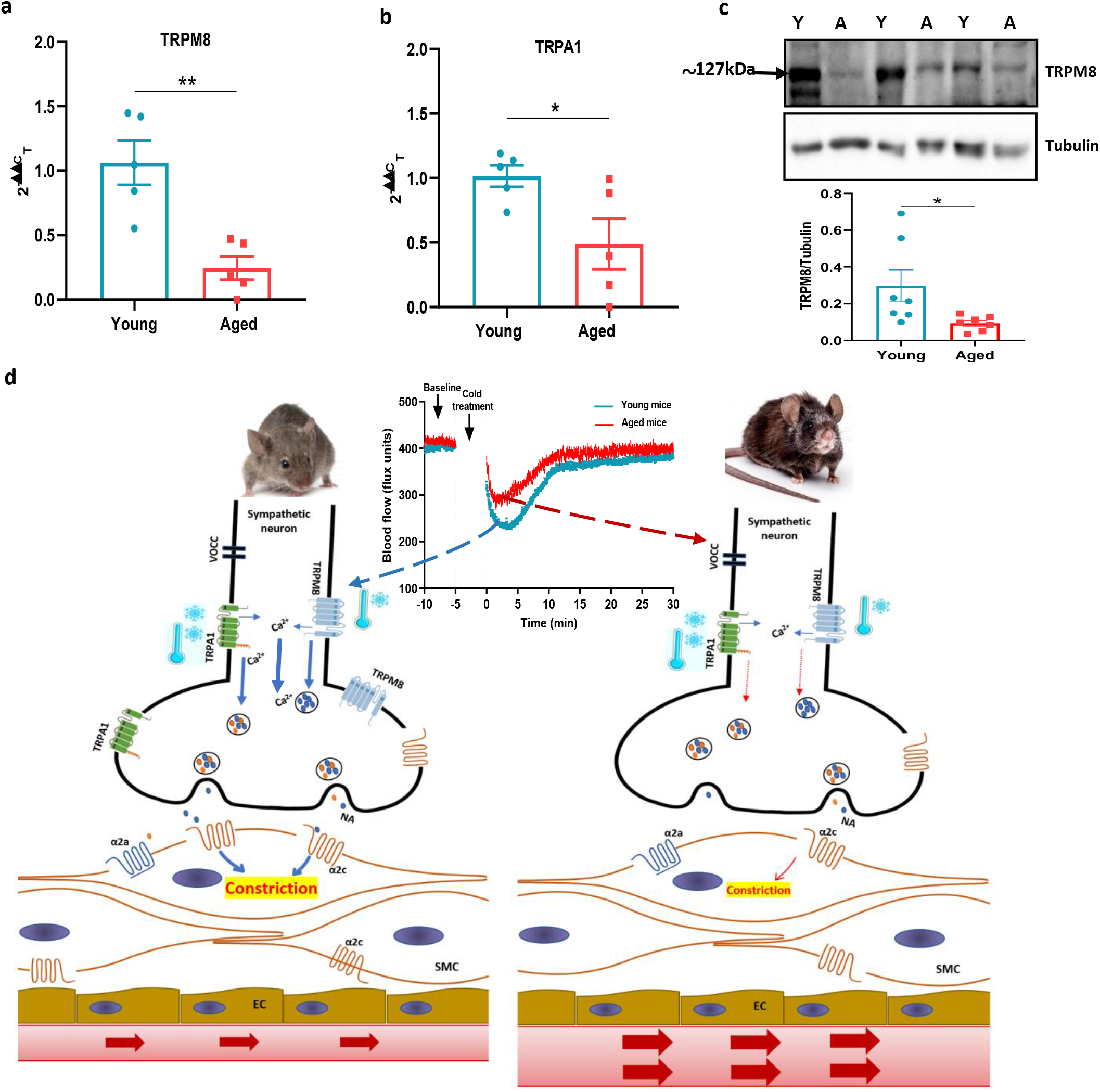
Sympathetic-sensory signalling and influence of ageing. **(a-b)** RT-PCR CT analysis shows the expression and fold change of TRPA1 and TRPM8 in young and aged sympathetic ganglia normalized to three housekeeping genes collected from the cervical and thoracic paravertebral region. **(c)** The western blot analysis of TRPM8 in sympathetic ganglia of young and aged mice. All results are shown as mean ± s.e.m. *p<0.05, **p<0.01. (Two-tailed Student’s t-test). **(d)** Proposed cold-induced vasoconstriction signalling pathway in young and aged mice. The local cold exposure produces rapid vasoconstriction which is significantly blunted in the aged mice (see blood flow graph at top centre). Cold water (4°C) exposure to hindpaw activates the cold receptors TRPA1 and TRPM8 in sympathetic nerves, which leads to increased intracellular calcium and release of NA. This signalling, however, is significantly downregulated in aged mice due to diminished expression of TRPA1/TRPM8 in sympathetic nerves. NA acts on the post-synaptic α_2c_ adrenergic receptors on smooth muscle cells to mediate vasoconstriction. However, α_2c_ adrenergic receptor are also significantly diminished in aged mice, which leads to reduced signalling. All of these factors contribute to an attenuated vascular cold response in aged mice compared to young mice. α_2c_ – alpha2c adrenoceptor, α_2a_ – α_2a_ adrenoceptor, VOCC- voltage operated calcium channel, NA – noradrenaline, Ca^2+^ - calcium, SMC-smooth muscle cell, EC – endothelial cell.

## Discussion

The role of TRPA1 and TRPM8 as cold-sensitive thermoreceptors is established (Story et al., 2003, Bautista et al., 2007, Karashima et al., 2009, Peier et al., 2002, McKemy, Neuhausser & Julius, 2002) and our research has demonstrated the essential role they play as vascular cold sensors (Aubdool et al., 2014, Pan et al., 2018). Much less was known about their activity in ageing, until this study. We provide a new insight into the changing roles of TRPA1 and TRPM8 in the vascular response to cold in ageing; the expression and activity of TRPM8 is significantly diminished, and to a lesser extent TRPA1 mediated signalling too.

The vascular response to the cold is a primary physiological response, which we have previously teased out the key mechanisms for, consisting of TRPA1/M8 initiated α_2c_-mediated sympathetic vasoconstriction followed by TRPA1/M8 mediated sensory vascular relaxation after localised cold exposure in the mouse paw. Here, we show that the response is functionally deficient in ageing as measured by two different laser blood flow measurement techniques. Both components of the cold-induced vascular response were impaired in ageing with blunted vasoconstriction that will lead to increased heat loss, and a slower rate of recovery that may lead to cold-induced injuries (Keatinge, 1957, Roustit et al., 2011, Herrick, 2005). We were surprized that the diminished response was observed with even moderate ageing (13-15 months old mice; equivalent to middle age in humans). However, the ageing nature of the mice was confirmed by the observation of increased expression of the established ageing markers p16 and p21 (Baker et al., 2016, Sharpless, Sherr, 2015, Hudgins et al., 2018). This finding is in keeping with the concept that although ageing-induced pathological conditions and frailty appear at a later age, the underlying physiological changes that manifest those conditions begin at a middle age of around 40 years old in humans. Indeed, this is when the brain volume and weight start to decline (Peters, 2006); cardiovascular functions begin to decline and elite athletes start to lose stamina (Pal et al., 2014, Mühlberg, Platt, 1999). We found that the baseline skin temperature was significantly higher in the aged mice than young mice, potentially due to greater heat loss from aged mice in keeping with the notion that it is harder to maintain core body temperature as ageing occurs. The tissue oxygen saturation was reduced during the vascular cold response but recovered substantially in the young compared to the aged mice. Overall, the findings show that the cold induced cutaneous vascular response is significantly diminished in ageing.

It is known that sensory modalities decline with ageing, but usually studies involve frail 24-month old mice. Indeed, in one of the only studies of TRP thermo-receptors in ageing, to our knowledge, authors investigated the changes in TRPM8 expressing neurons in cornea with ageing and its relevance to dry eye disease (Alcalde et al., 2018). Here in our study, we delineate that activity and expression of one of the prominent sensory cold channels (TRPM8) diminishes with moderate ageing; relevant to the impaired vascular response to cold that we have observed. We designed experiments to evaluate the ability of the mouse to sense cold using a cold plate behavioural assay at innocuous cool (TRPM8 range) and noxious cold (TRPA1 range) temperatures. We observed a delayed latency to the cold response in moderately aged mice compared to young mice at three different cold temperatures, 4°C, 10°C, and 20°C implying impaired sensory signalling of TRPM8 and TRPA1 receptors. Importantly, the longest delayed latency was observed at 20°C, which falls under the TRPM8 activation range implying that with ageing TRPM8 signalling deteriorates more, relative to that of TRPA1.

Using combined selective antagonists of TRPA1 and TRPM8, we show that blocking both receptors inhibits the cold-induced cutaneous vascular response in young mice, as expected.

Intriguingly, the same treatment produced a relatively stronger inhibitory response in the aged mice. This implies that with ageing the cold signalling relies profoundly more on cold TRP channels. These results may also indicate that at younger ages other protein/s besides TRPA1 and TRPM8 play a crucial part in the cold signalling, but these activities begin to decline with ageing. This includes the α_2c_ receptor as discussed below, although a range of other candidates have also been proposed (Zimmermann et al., 2011, Noël et al., 2009, Luiz et al., 2019, Gong et al., 2019). Importantly, when we investigated the effect of the TRPA1 antagonist alone, we observed a very similar inhibitory profile to that of the combination of TRPA1 and TRPM8 antagonists. However, the TRPM8 antagonist treatment alone, was effective in the young mice, but not in the aged mice. This provided key evidence that the activity of TRPM8 especially, is diminished in ageing. TRPM8 was discovered as a sensory receptor expressed in DRG and trigeminal ganglia (TG) that is activated by cool temperatures (<28°C) (McKemy, Neuhausser & Julius, 2002); although the link with TRPA1 containing CGRP fibres is less well defined (Hondoh et al., 2010, Kobayashi et al., 2005). Since then, various reports have suggested that TRPM8 is expressed in a wider range of tissues and is involved in multiple physiological functions including thermoregulation (Moraes et al., 2017, Hirai et al., 2018, Yang et al., 2006). Indeed, it is established that the deletion/antagonism of TRPM8 increases heat loss and reduces core body temperature (Almeida et al., 2012, Reimúndez et al., 2018).

By this stage we had revealed a reduced expression and activity of TRPM8 in the vascular cold response and cold sensing. However, we have previously defined TRPA1 as an essential vascular sensor to cold, playing a major role in the cold induced vascular response alongside TRPM8 (Aubdool et al., 2014). Therefore, it was surprising to observe that expression of TRPA1, unlike TRPM8, did not diminish in the DRG with ageing, especially as the cold-sensing data from the cold plate at noxious cold temperatures revealed that the response is also impaired at noxious temperatures in ageing. To learn more, we studied the ability of the TRPA1 agonist cinnamaldehyde (CA) to increase cutaneous blood flow; as TRPA1 is localised to CGRP^+^ sensory nerves (Aubdool et al., 2016). CA-induced vasodilation was significantly impaired in the aged mice compared to young, supporting the concept of impaired functional TRPA1 vascular responses in ageing. We observed a similar significantly blunted response with the TRPM8 agonist menthol in aged mice, although TRPM8 localisation to sensory nerves is more limited than that of TRPA1 (Kobayashi et al., 2005). This led us to question whether activity of all TRP receptors deteriorates with ageing, through investigating the non-cold sensing TRPV1 agonist capsaicin which activates predominantly CGRP^+^ C-fibres (Story et al., 2003, Sharrad et al., 2015). In contrast to menthol and CA, capsaicin caused a similar level of increased blood flow in all mice regardless of age indicating TRPV1 activity does not deteriorate with ageing, and supporting our behaviour data at 30°C, which falls outside TRPA1 and TRPM8 activation ranges. These results suggest that only the signalling of cold TRP receptors; TRPA1 and TRPM8 is impaired with ageing.

The cold-induced vasoconstriction phase is mediated by sympathetic drive comprising of noradrenergic nerves and this signalling has been shown to decline with ageing (Degroot, Kenney, 2007, Frank et al., 2000, Greaney, Alexander & Kenney, 2015). Thus, we aimed to elucidate the sympathetic signalling in young and aged mice, which we began by investigating the effect of exogenous agonist NA. NA administered locally to the paw evoked cutaneous vasoconstriction in young mice that was significantly blunted in the aged mice, suggesting that NA-mediated response diminishes in aged mice. Nonetheless, NA is a non-selective agonist for all adrenoceptors, but peripheral cutaneous vasoconstriction is mediated via α adrenergic receptors (Drew, Whiting, 1979), with cold specific vasoconstriction primarily mediated via α_2c_ adrenoceptors subtype (Bailey et al., 2004, Honda et al., 2007). Therefore, we used the selective α_2_ agonist medetomidine which induced vasoconstriction that was also significantly blunted in the aged mice. The result suggests that either α_2c_ receptor sensitivity declines with ageing (Thompson, Holowatz & Kenney, 2005) or α_2c_ receptor number reduces with ageing which has been suggested to occur in ageing human saphenous vein (Hyland, Docherty, 1985). In our study, we found a significant reduction in the expression of α_2c_ adrenoceptors. We also investigated whether the level of NA or its synthesis was impaired in ageing and observed no difference in the level of tyrosine hydroxylase (including the active form of phosphorylated tyrosine hydroxylase), the enzyme involved in the rate limiting synthesis of NA production. This indicates that NA synthesis is not affected in ageing.

The cold-induced vascular response is perceived as a reflex where peripheral sensory nerves sense the cold stimulus and send information to the central nervous system (CNS). In turn, the CNS processes the information and produces an appropriate response via activation of sympathetic nerves to cause vasoconstriction in skin (Chotani et al., 2000). Classically, it is established that sensory receptors TRPA1 and TRPM8 that sense cold reside in sensory nerves and alpha-adrenergic receptors reside in sympathetic nerves and smooth muscle cells to modulate vasoconstriction. However, we have previously shown that the cold-induced vasoconstrictor response occurs when the sensory C-fibre component is removed with resiniferatoxin treatment (Aubdool et al., 2014). This clear result raises the possibility that the cold sensitive proteins TRPA1 and TRPM8 may be expressed in other tissues besides sensory nerves and modulate the vascular tone, as suggested to be the case in some organs (Earley, 2012, Johnson et al., 2005, Yang et al., 2006). The sensory nerves and sympathetic nerves are known to have close proximation around blood vessels and have a reciprocal trophic influence (Terenghi et al., 1986). Thus, we questioned whether cold TRP channels were expressed on sympathetic nerves to directly modulate the vascular tone. We found that both TRPA1 and TRPM8 are expressed in the sympathetic ganglia (SG) collected from the cervical and thoracic paravertebral regions, in keeping with previous studies (Smith et al., 2004), but debated. Furthermore, the expression of both receptors were significantly diminished in the SG collected from the aged mice compared to young mice. Collectively, these findings suggest that cold stimuli activate TRPA1 and TRPM8 channels on sympathetic nerves, which induces calcium-dependent release of vesicles containing NA into the synaptic cleft where they activate the α_2c_ adrenoceptor on smooth muscle cells to mediate vasoconstriction (Fig 7d). This signalling cascade has been shown in PC12 cells, which is regularly used as *in-vitro* model for sympathetic neurons (Smith et al., 2004, Peixoto-Neves, Soni & Adebiyi, 2018, Yoshimura, Nakagawa & Endo, 2016). It indicates a potential for sympathetic-sensory interactive signalling in skin, which weakens as ageing progresses in turn affecting the sensitivity of the vascular response to cold.

To conclude, we have revealed that the cold induced defensive responses decline with ageing. There is an impairment in the sympathetic vasoconstrictor pathway concomitant with a functional deterioration and molecular loss of TRPM8 and TRPA1 signalling as well as diminished α_2c_ adrenergic receptor expression and activity. We consider that the finding of diminished TRPM8 expression with ageing is indicative of a major influence of this channel that would lead to the impaired cold induced vascular response in ageing.

## Methods

### Animals

Female CD1 mice used in this study were either bred in the Biological Services Unit, King’s College London or purchased from Charles River (Kent, UK). The animals were housed in a climatically controlled environment with an ambient temperature of 22°C, including a 12-hour light/dark (7am-7pm) cycle with free access to drinking water and standard chow *ad libitum*. Young mice were 2-3 months old and aged mice were 13-16 months old. All experiments were performed according to the Animal Care and Ethics committee at King’s College London, in addition to the regulations set by the UK home office Animals (Scientific Procedures) act 1986. Experiments using animals were designed and reported in line with the ARRIVE guidelines, which form the NC3Rs initiative. Animals were randomly assigned to different groups and the investigator was blinded to drug treatments and where possible to the age of the animals.

### Cutaneous blood flow measurement by full-field laser perfusion imager

Mice were terminally anaesthetized with i.p. injection of ketamine (75 mg kg^−1^) and medetomidine (1 mg kg^−1^). Wherever possible to comply with the NC3Rs reduction guidelines, experiments were designed using recovery anaesthesia, which was either s.c. 150 mg kg^−1^ ketamine and 4.25 mg kg^−1^ xylazine, or isoflurane gas. 5% isoflurane (in oxygen) was used to induce anaesthesia, which was followed by 2% for maintenance during the experimental procedure. Full-field Laser Perfusion Imager (FLPI, Moor Instruments, UK) was used to measure blood flow in the hind paw or the ear of the mice. The mice were placed in a ventral position on a heating mat to maintain core body temperature at 37°C during blood flow measurement. For the cold-induced blood flow measurement in hindpaw, after anaesthesia, the blood flow was measured on the plantar surface of hindpaw for 5 min as baseline measurement. Then, the ipsilateral hindpaw at the level between tibia and calcaneus was immersed in cold water (4°C for 5 min) for cold exposure. After the cold treatment, mice were placed back on the heating mat (37°C) to measure blood flow recovery for 30 min. The FLPI uses laser light to produce speckle pattern that gets interfered by blood flow which is measured as arbitrary flux units (X10^3^ flux units). For the agonist-induced blood flow measurement in ear, after anaesthesia, the blood flow was measured for 5 min as baseline recording. 10μl of either cinnamaldehyde (10%), menthol (10%), or capsaicin (10%) was topically applied to both sides of the ipsilateral ear and 10μl vehicle solution (10% DMSO in ethanol) was applied to the contralateral ear. Then blood flow was measured for 30 min after cinnamaldehyde and capsaicin treatment and for 10 min after menthol treatment. All treatments produced a gradual increase in blood flow. For NA/medetomidine-induced blood flow measurement in hindpaw, after anaesthesia, blood flow was measured on the plantar surface for 5 min as baseline recording. Intraplantar injection of NA/medetomidine (1.25ng/ul) was performed and blood flow was measured for 15 min.

### Cutaneous blood flow, temperature and oxygen saturation measurement by laser Doppler techniques

A probe connected to the moorVMS-LDF (Laser Doppler Perfusion and Temperature Monitor) and moorVMS-OXY (Tissue Oxygen and Temperature Monitor) (Moor Instruments) was used to simultaneously measure blood flow, temperature and tissue oxygen saturation in a small, localized area (~5mm diameter) on the plantar surface of the ipsilateral hind paw (central region immediately adjacent to the digits). The probe was held on a retort stand clamp 1mm above the skin surface. After inducing anaesthesia with i.p. injection of ketamine (75 mg kg^−1^) and medetomidine (1 mg kg^−1^), the blood flow was measured (baseline recording) on the plantar surface central area immediately adjacent to the digits for 5 min. The ipsilateral hindpaw was immersed in cold water (4°C for 5 min) at the level between tibia and calcaneus. After the cold treatment, mice were placed back on the heating mat (37°C) to record all measurements during the recovery period for 30 min. The blood flow was measured using doppler technique and expressed in arbitrary flux units, and tissue oxygen saturation was measured using white light spectroscopy method.

### Drugs and reagents

The TRPA1 antagonist A967079 ((1E,3E)-1-(4-Fluorophenyl)-2-methyl-1-pentene-3-one oxime) (Alomone Labs, # A-225) was dissolved in 10% DMSO, 10% Tween-80 in saline. The TRPM8 antagonist AMTB (N-(3-aminopropyl)-2-[(3-methylphenyl) methyl] oxy-N-(2-thienylmethyl) benzamide hydrochloride salt) (Alomone Labs, #A-305) was dissolved in 10% DMSO in saline. Both antagonists were administered i.p. 30 min before the cold treatment. Cinnamaldehyde (Sigma Aldrich, #W228613, >95% purity), menthol (Alfa Aesar, #A18098, 98% purity), capsaicin (Sigma Aldrich, #M2028, >95% purity) were prepared with 10% DMSO in ethanol solution. 1.25ng/μl NA (Sigma) and 1.25ng/μl medetomidine (Orion Pharma) were administered with intraplantar injection in 20μl saline.

### Behavioural testing using the cold plate

The nociceptive cold sensitivity response of mice was tested using a hot/cold thermal plate (Ugo Basile 35100). A quick temperature non-contact infrared thermometer (Linear labs) was used to confirm the set temperature of the plate before each experiment. Prior to the experiments, mice were acclimatised to the room for 30 min for 3 days, and the thermal plate by individually placing them on the plate at room temperature for 2 min on each of the 3 days. At the start of the experiment, the plate was set to the chosen temperature (4°C, 10°C, 20°C and 30°C) and each mouse was placed individually onto the plate in turn. The cold response was detected as either paw licking or paw withdrawal/jumping and the total number of responses observed within 1 min (for 4°C and 10°C) and 5 min (20°C and 30°C) were tallied. Each temperature was repeated twice on different days to obtain an average which was used to plot the final graph.

### Quantitative polymerase chain reaction

Real time PCR (RT-qPCR) was used to quantify changes in mRNA collected from pooled dorsal root ganglia (DRG), brown adipose tissue (BAT) and sympathetic ganglia collected from thoracic paravertebral region. The total RNA was isolated and purified according to manufacturer’s instructions using the RNeasy Micro Kit (Qiagen, #74004). The RNA concentration and absorbance ratio (A260/280 and A260/230) were measured using Nanodrop 2000 spectrophotometer. 500ng of purified RNA was reverse transcribed using SuperScript ViLO cDNA synthesis kit (Thermo fisher scientific, #11754050). qPCR was performed with 10ng of cDNA using PowerUp SYBR Green master mix kit (Thermo fisher Scientific, #A25780) in 7900HT Real-Time PCR machine (Applied Biosystems, USA). All primers (Supplementary table 1) were designed using Primer-BLAST software (NCBI) according to MIQE guidelines and checked on the primer stat website. (http://www.bioinformatics.org/sms2/pcr_primer_stats.html). The melting curve analysis was performed after reactions to confirm specificity of the primers. The analysis was performed using delta delta CT method and expressed as fold change normalized to the average of three housekeeping genes.

### Western blotting

The western blotting analysis was performed as previously described (Aubdool et al., 2014). The tissue was lysed with SDS lysis buffer which was made up with inhibitors of both phosphatases and proteases (1 tablet per 10ml, #4693159001 + #4906845001, Sig-ma-Aldrich). The tissue was then homogenised using a tissue lyser (Qiagen, #85300). The protein concentration was determined using the Bradford protein dye binding assay (#5000113 + #5000114, Bio-Rad). 50μg of protein was separated by electrophoresis in an SDS-polyacrylamide gel which was then transferred using the semi dry method, onto PVDF membranes. The membrane was incubated in a blocking buffer made up of 5% BSA in Phosphate-buffered saline- tween (PBS-T) (0.1% Tween). The membrane was blocked for 1 hr in RT except for TRPM8 which was blocked for 2.5 hr as per manufacturer’s instruction. The membranes with primary antibodies were incubated overnight at 4°C. Following the washing step with PBS-T, the membranes were probed with secondary antibody (Horseradish peroxidase conjugated) (1:2000 dilution, #AP132P Sigma) for 1 hr at RT. The enhanced chemiluminescence (ECL, Piercenet) was used for visual development of the membranes inside a gel doc system. Bands were normalised to housekeeping genes α-tubulin (1:2000, #MAB1864, Merck Millipore), GAPDH (1:2000, #PA1987, Thermofisher) and β-actin (1:2000, #A5441, Sigma Aldrich). Quantitative western blot analysis was performed using Image J (NIH, USA). The primary antibodies were made in 3% PBST solution at 1:500 dilution for TRPM8 (Alomone Labs #ACC049), 1:1000 dilution for α_2c_ adrenergic receptor (Bio-Techne #NB100-93554), phospho TH (Bio-Techne #NB300-173), total TH (Bio-Techne #NB300-109), p21 (Santa Cruz #sc-6246).

### Experimental design and data analysis

The majority of the experiments conducted in this study consisted of two groups (young/aged) or four groups with drug treatments (young/aged and vehicle/drug), therefore the power analysis from our lab(Aubdool et al., 2014) with a power of 80% (0.8) for a confidence of 5% (0.05) recommended n=8, which was strictly adhered to where possible. The order of the mice (young or aged) and treatments (vehicle/drug) received were randomised during experimental protocols. Data was analysed using either two-tailed Student’s t-tests or two-way ANOVA followed by Tukey’s *post hoc* test. All column data are plotted as dot plots to show variability and n numbers for each data set. All data are expressed as mean ± SEM. p<0.05 was considered to represent a significant difference. GraphPad Prism (version 8) was used as statistics software for analysis.

## Acknowledgements

This work was primarily funded by BBSRC (BB/P005616/1). It was also supported in part by Versus Arthritis (ARUK21524) and British Heart Foundation (BHF-FS/19/42/34527 and PG/12/34/29557).

## Author contributions

DT and SDB designed the research. DT, JV, BB, FA, SL and SN carried out research. DT, BB and SL performed data analysis. XK helped with blinding and data analysis. DT, BB and SDB drafted the manuscript and all authors contributed to finalizing the manuscript.

## SUPPLEMENTARY INFORMATION

**Supplementary Figure 1:**
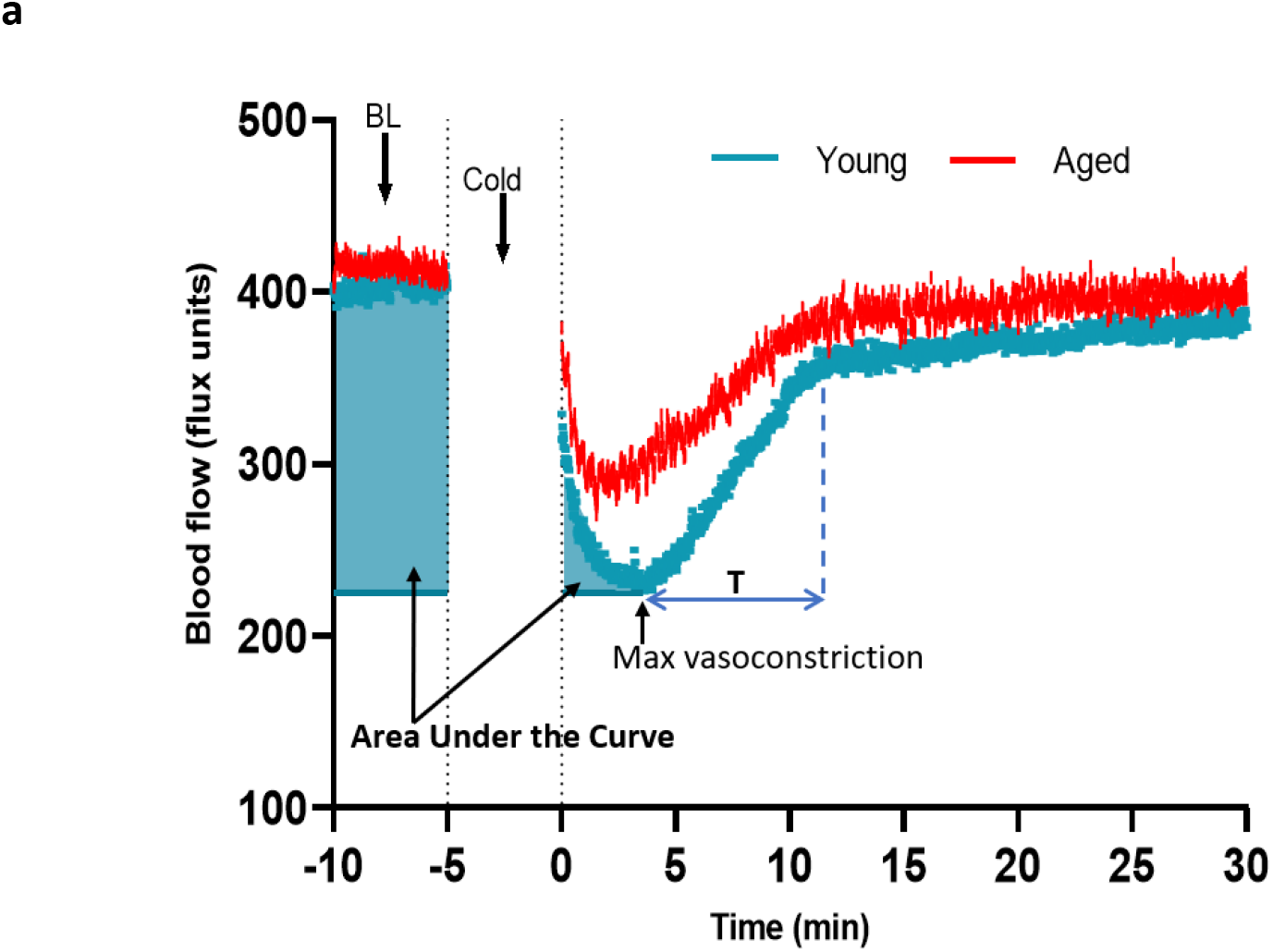
Analysis of cold-induced blood flow in the mouse paw. **(a)** The blood flow graph from cold treatment was used to calculate the maximum vasoconstriction, area under the curve (AUC) analysis of vasoconstriction and recovery of blood flow from cold treatment. It was not possible to measure blood flow whilst the paw was being cooled. The highlighted areas in blue shows the area of the graph from start of the 5 min baseline (BL) until peak vasoconstriction that was used to calculate the AUC analysis to detail the magnitude of the vasoconstrictor event. The time of immediate blood flow recovery (T) was calculated by measuring the time immediately after maximum vasoconstriction until it started to plateau back to baseline level (as shown by blue lines). (BL = Baseline). The maximum change of blood flow (%) was calculated using the equation below:

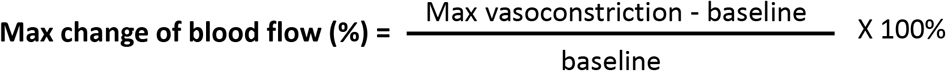

**Supplementary Figure 2:**
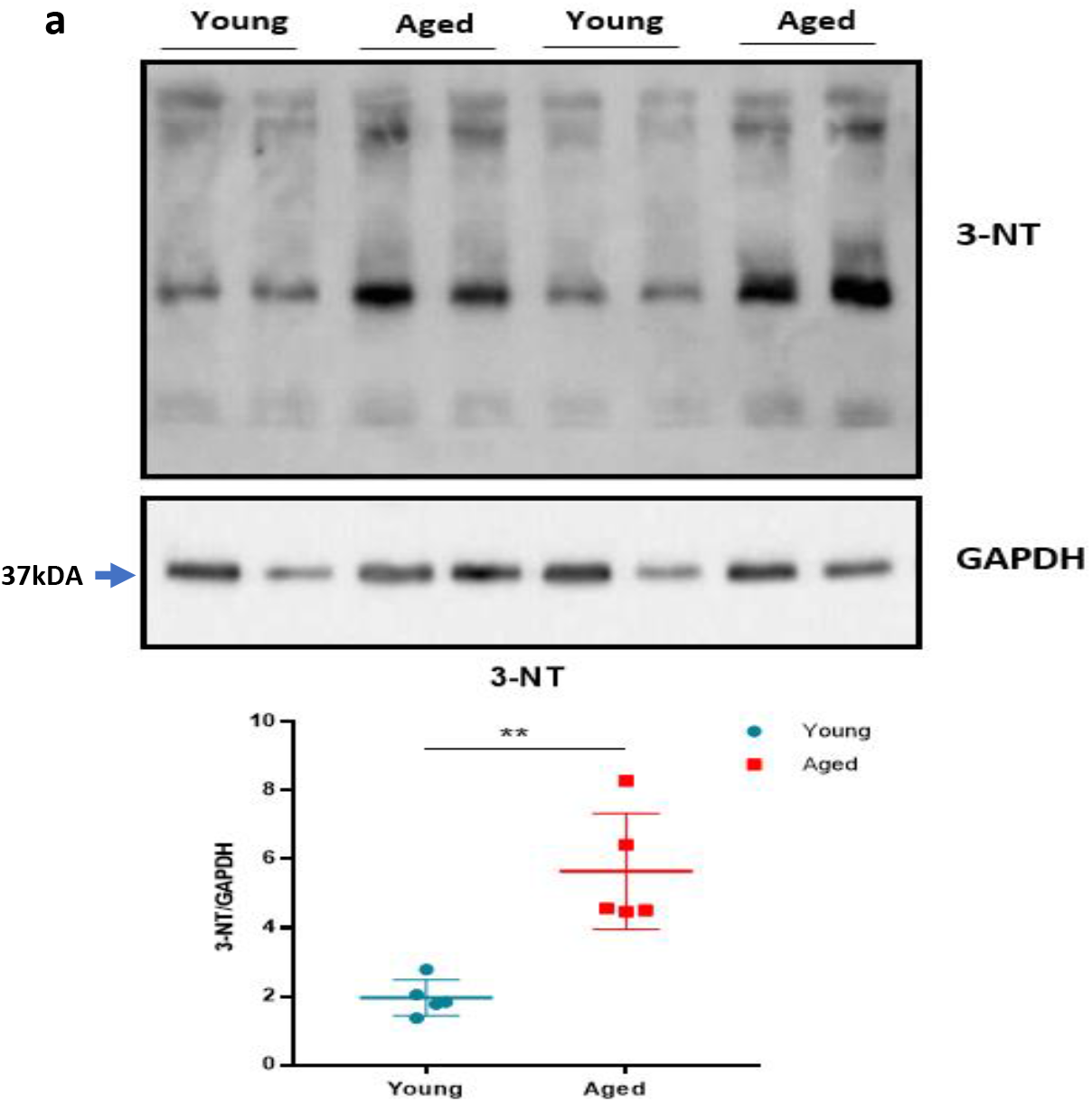
Oxidative stress with ageing. **(a)** The representative western blot analysis of 3-nitrotyrosine (3-NT), which is a biomarker of oxidative stress produced by reactive nitrogen species, in hind paw skin of naïve young and aged mice. The densitometric analysis is shown normalized to GAPDH housekeeping gene. Results are shown as mean ± s.e.m. (n=5) **p<0.01. (Two-tailed Student’s t-test).

**Supplementary Figure 3:**
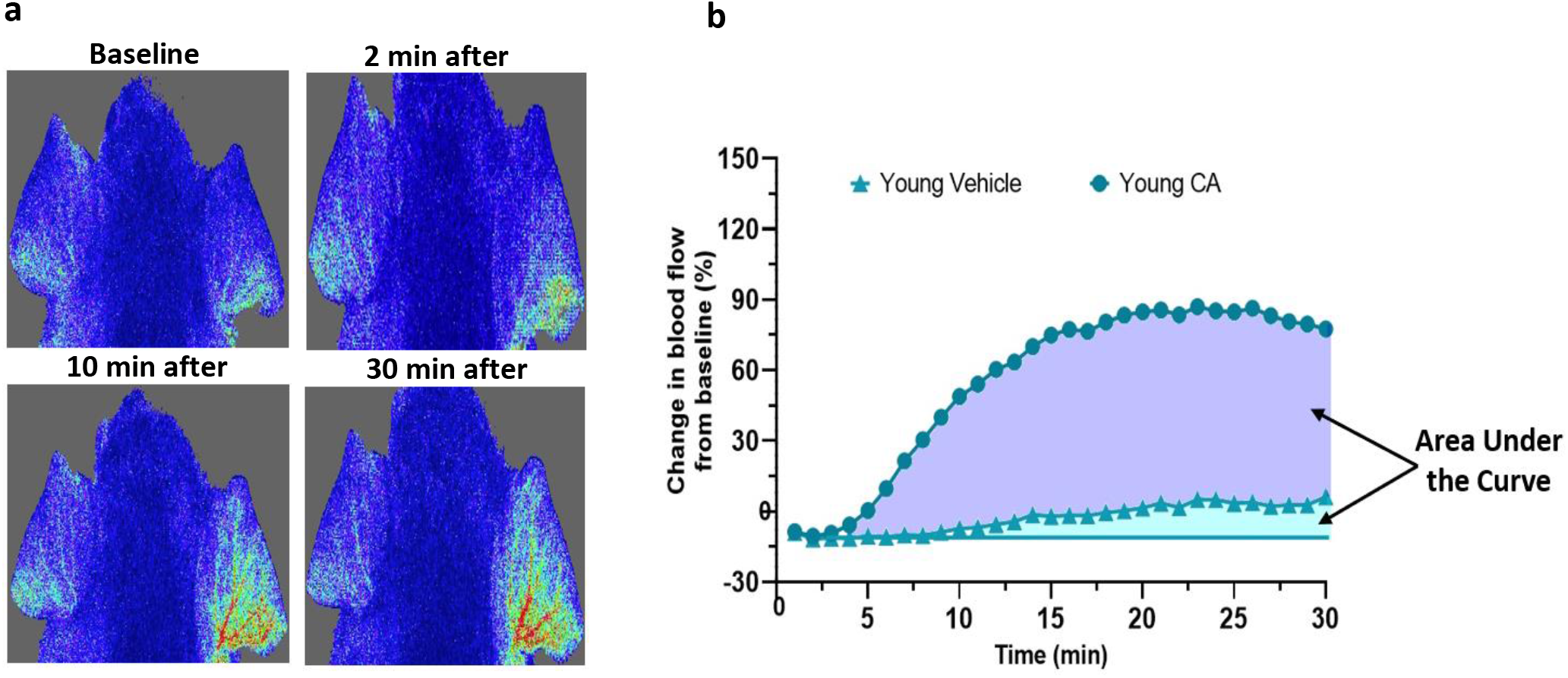
Agonist induced blood flow response in mouse ear. **(a)** The representative image from FLPI shows the blood flow response induced by topical application of vehicle (left ear) and 10% cinnamaldehyde (right ear) in the anaesthetised mouse. **(b)** The representative blood flow response graph illustrates the area used for the area under the curve analysis for vehicle (light blue) and cinnamaldehyde (purple).

**Supplementary Figure 4:**
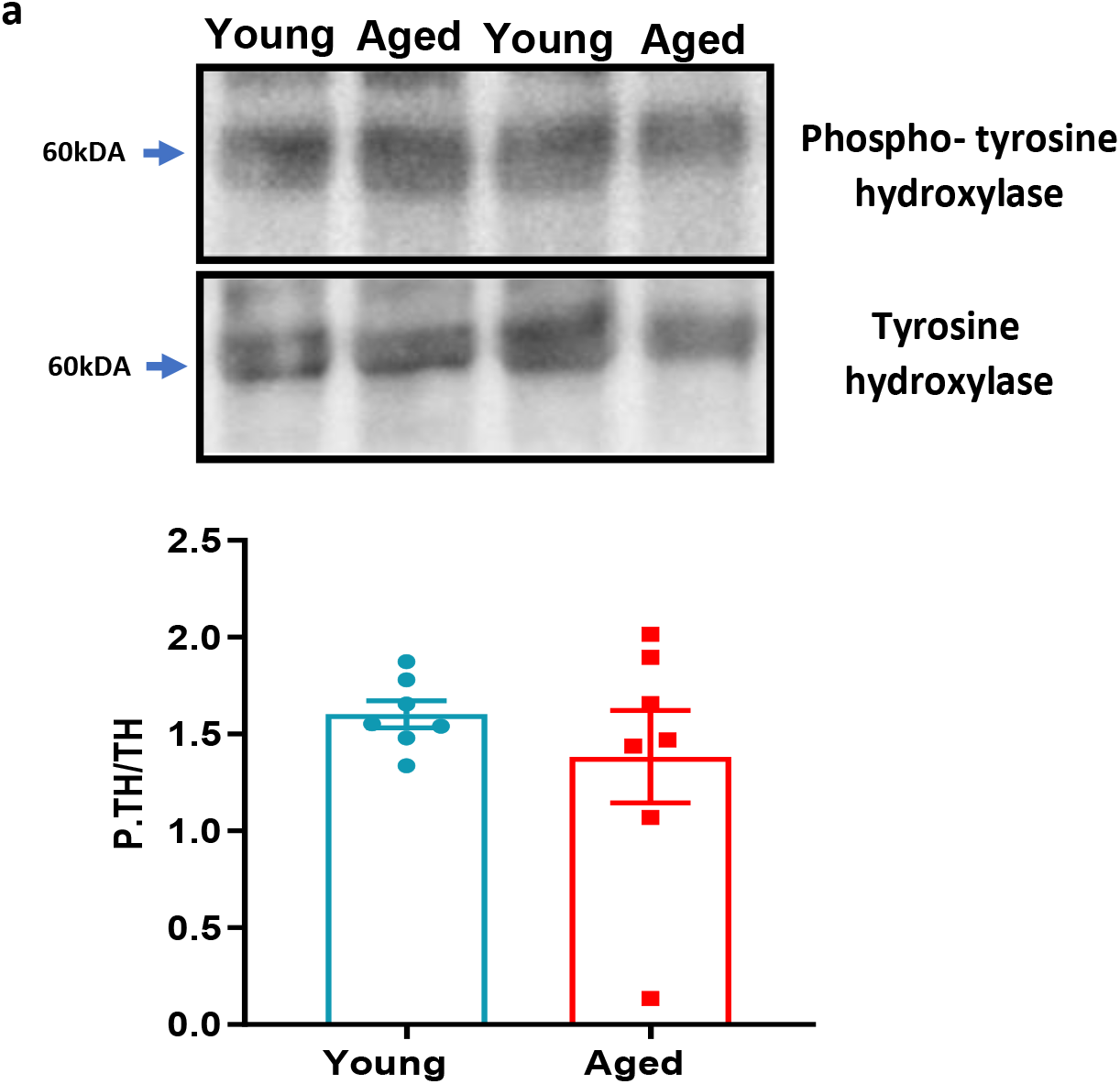
Phospho-tyrosine hydroxylase with ageing. **(a)** The representative western blot analysis of phospho-tyrosine hydroxylase in hind paw skin of naïve young and aged mice. The densitometric analysis is normalized to total tyrosine hydroxylase. Results are shown as mean ± s.e.m. (n=7) (Two-tailed Student’s t-test).

**Supplementary Figure 5.**
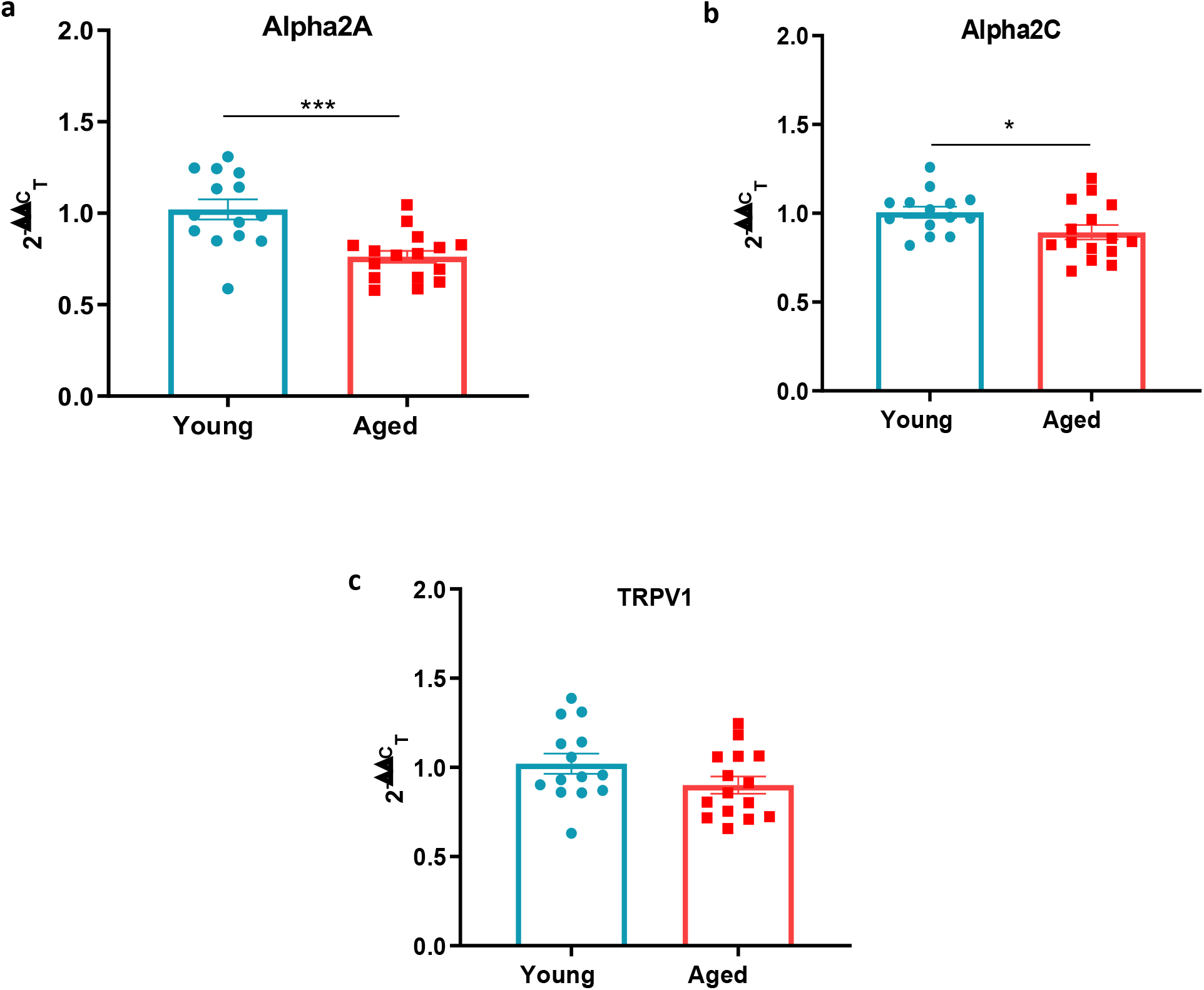
Sympathetic-sensory signalling in young and aged mice. **(a-c)** RT-PCR CT analysis shows fold change of α_2a_ adrenoceptor, α_2c_ adrenoceptor and TRPV1 in young and aged mice normalized to three housekeeping genes in dorsal root ganglia (DRG). Results are shown as mean ± s.e.m. (n=14-16) *p<0.05, ***p<0.001. (Two-tailed Student’s t-test).

**Supplementary Figure 6.**
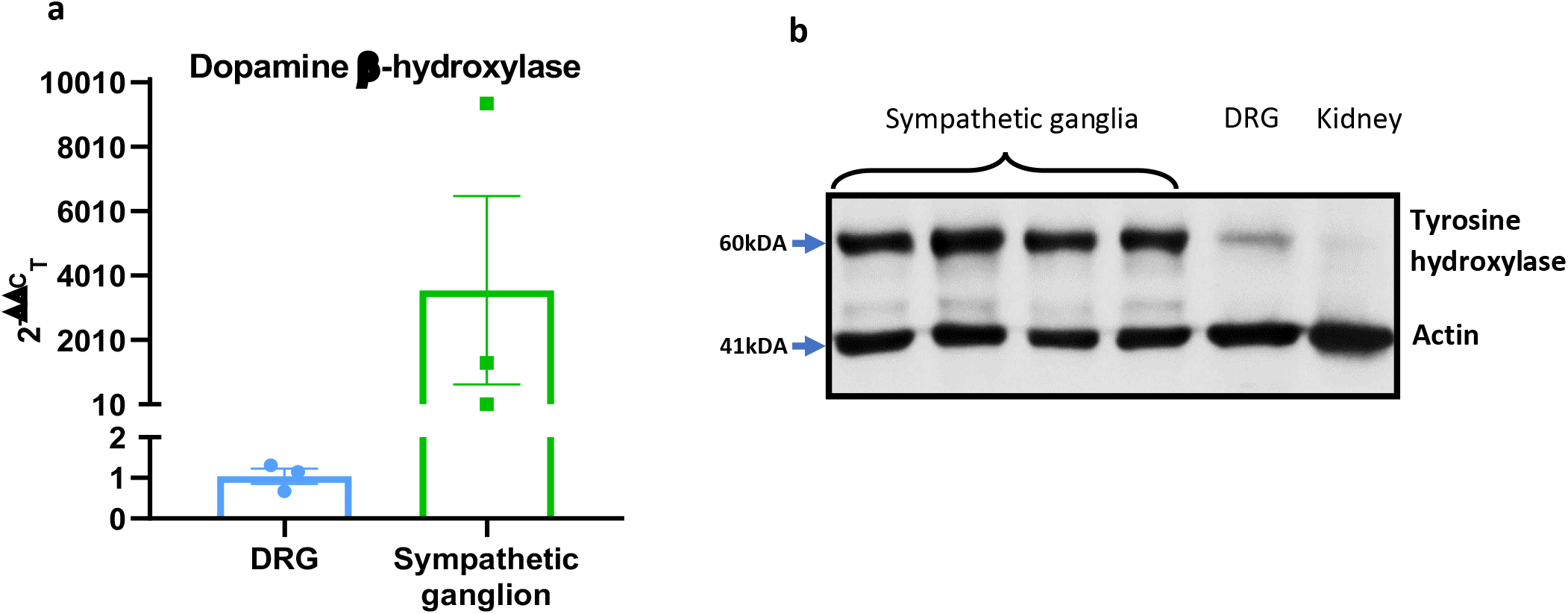
Characterisation of sympathetic markers. **(a)** RT-PCR CT analysis shows fold change of the sympathetic nerve marker dopamine β-hydroxylase (Dbh) in DRG and sympathetic ganglia normalized to three housekeeping genes. **(b)** The western blot analysis of the sympathetic nerve marker tyrosine hydroxylase (TH) in sympathetic ganglia, DRG and kidney. Results are shown as mean ± s.e.m. (n=3).

**Supplementary Table 1.**
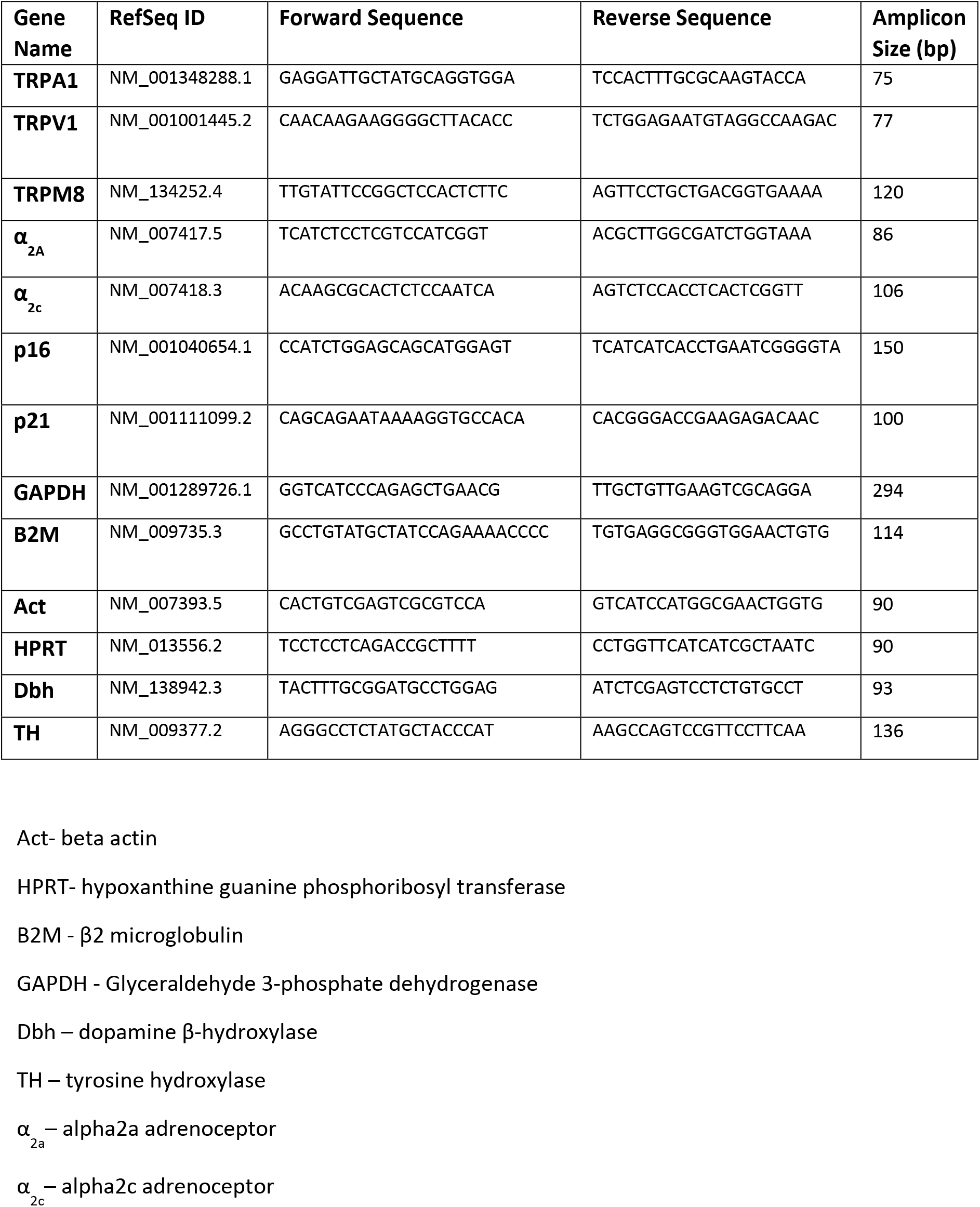
List of primer sequences

